# Mapping the spatiotemporal dynamics of *de novo* protein synthesis during long-term memory formation

**DOI:** 10.1101/2025.04.17.649250

**Authors:** Harrison T. Evans, Drew Adler, Maria Clara Selles, Srinidhi V. Kalavai, Astra Yu, Victor Wu, Eléa Denil, Ela N. Golhan, Ambika Polavarapu, Emma Balamoti, Moses V. Chao, Robert C. Froemke, Eric Klann

## Abstract

The formation of new associative long-term memory (LTM) following Pavlovian conditioning is dependent upon multiple, temporally distinct windows of mRNA translation. Current methods lack the temporal specificity to robustly characterize the dynamics of protein synthesis throughout the rodent brain following conditioning. Here we resolve these technological limitations and demonstrate that in awake mice, the retro-orbital (RO) injection of azidohomoalanine (AHA) enables the labelling and subsequent visualization of the brain *de novo* proteome, with labelling periods as short as 30 minutes. Combining this advancement in *de novo* proteomic labelling with tissue clearing, we identified brain region, cell-type, and neuronal sub-population specific changes in *de novo* protein synthesis in mice following an auditory threat conditioning paradigm. This approach also allowed us to track the changes in *de novo* protein synthesis over time, revealing that conditioning-induced changes in mRNA translation exhibit remarkable temporal specificity in brain regions such as the somatosensory cortex. Taken together, our findings highlight how this novel labelling technique can be used to map the highly intricate temporal and spatial dynamics of mRNA translation after behavioral conditioning.

## Introduction

The formation of new long-term memories (LTM) is reliant upon mRNA translation. This was first established in 1963 by Flexner and colleagues who showed that intracerebral injection of puromycin, an antibiotic that inhibits protein synthesis, impaired avoidance discrimination memory in mice (Flexner *et al*, 1963). Since then, numerous studies have utilized pharmacological, and more recently, chemogenetic techniques to identify brain regions and cell types in which protein synthesis is required for the consolidation of different forms of long-term memory (Lima *et al*, 2009; Pedroza-Llinás *et al*, 2009; Ozawa *et al*, 2017; Shrestha *et al*, 2020a; Moncada & Viola, 2007).

Taken together, these previous findings led researchers to hypothesize that during memory consolidation, multiple spatially and temporally distinct windows of protein synthesis enable the formation of the neuronal circuits that encode memory (Shrestha & Klann, 2022). Indeed, many of the proposed cellular correlates of memory, such as long-term potentiation and long-term depression, have been demonstrated to require *de novo* protein synthesis (Younts *et al*, 2016; Fonseca *et al*, 2006). Furthermore, several neurodevelopmental and neurodegenerative diseases in which memory is impaired also exhibit perturbations in various aspects of translational control, resulting in either exaggerated or deficient *de novo* protein synthesis (Mohamed & Klann, 2023; Evans *et al*, 2019; Elder *et al*, 2021; Oliveira *et al*, 2021; Evans *et al*, 2021b; Koren *et al*, 2019). Thus, it is clear that mRNA translation plays a causal and critical role in the formation of new long-term memories.

Given the requirement of *de novo* protein synthesis for memory consolidation, identifying which proteins are newly synthesized, in which brain regions and cell types these proteins are synthesized, and during which stages of consolidation this synthesis occurs, is vital to understanding the spatial-temporal dynamics of memory. Unfortunately, technical limitations have prevented researchers from visualizing *de novo* protein synthesis during memory formation in the rodent brain.

The current gold-standard for labelling, visualizing, and identifying newly synthesized proteins is non-canonical amino acid (NCAA) tagging (Evans *et al*, 2021a) where newly synthesized proteins are labeled with azide-bearing NCAAs such as azidohomoalanine (AHA), which incorporates into the nascent polypeptide chain in the place of methionine (Fig. 1A). Unlike other *de novo* proteomic techniques, such as puromycin labeling, AHA labeling has minimal effects on protein structure or function (McClatchy *et al*, 2020). The azide group of AHA can be used to covalently bond labeled newly synthesized proteins to a variety of tags via the azide-alkyne cycloaddition. As a result, newly synthesized proteins can be either visualized via FUNCAT (fluorescent non-canonical amino acid tagging) or isolated using BONCAT (bio-orthogonal non-canonical amino acid tagging) and subsequently identified via mass spectrometry (Carlisle *et al*, 2023; Dieterich *et al*, 2006).

**Figure 1:**
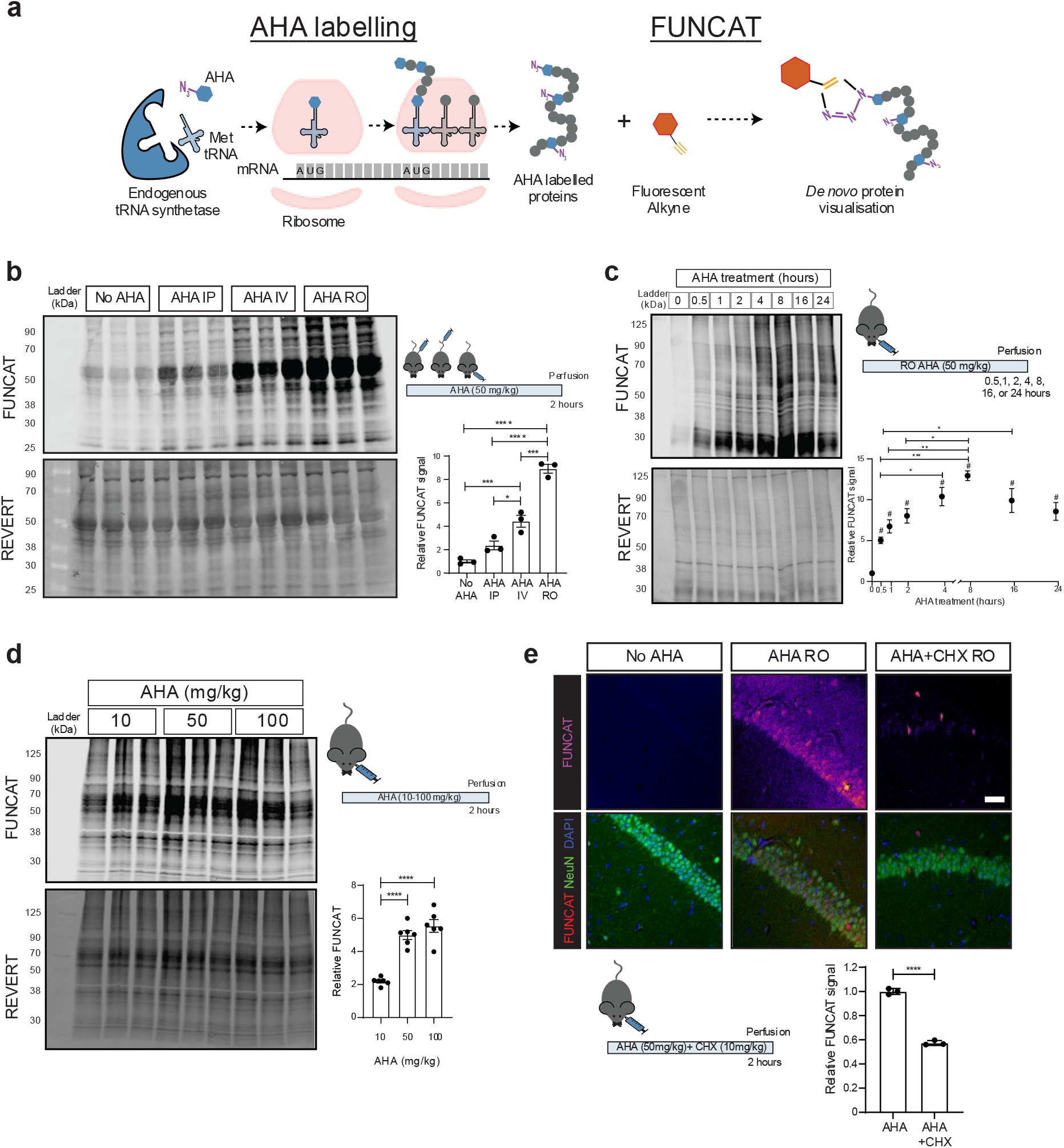
Retro-orbital injection of AHA allows for rapid labelling of the brain de novo proteome. **(A)** Newly synthesized proteins can be labeled by the methionine surrogate AHA and then be covalently bonded to a fluorescent tag using FUNCAT. **(B)** Retro-orbital delivery of AHA results in significantly higher brain *de novo* proteome labelling after 2 hours of treatment compared to i.p. and i.v. (intra tail-vein) injection as measured by FUNCAT-WB (one-way ANOVA, Tukey’s MCT, n=3 animals). **(C)** AHA labeling of *de novo* synthesized proteins can be detected as little as 30 minutes post retro-orbital (R.O.) injection, with FUNCAT signal peaking 8 hours post injection (one-way ANOVA, Tukey’s MCT, n=3 animals). **(D)** Treating mice with dosage of AHA higher than 50 mg/kg does not increase *de novo* proteomic labelling, as measured via FUNCAT-WB (one-way ANOVA, Tukey’s MCT, n=6 animals). **(E)** Co-injection of the protein synthesis inhibitor cycloheximide (CHX) significantly reduces AHA labeling as measured by FUNCAT-IHC (Students T.Test, n=3 animals, ≥3 sections per animal). Scale bar = 50μm. Error bars = S.E.M. *= p≤ 0.05, **= p≤ 0.01, ***= p≤ 0.001, ****= p≤ 0.001

Labeling newly synthesized proteins with NCAAs offers several distinct advantages over other *de novo* proteomic analysis techniques. Unlike techniques that rely upon isotopically labeled amino acids such as SILAC (stable isotype labeling with amino acids in cell culture), newly synthesized proteins labelled with NCAAs can be either visualized or purified from the rest of the proteome (Hinz *et al*, 2013). In addition, the labelling of newly synthesized proteins with NCAAs has minimal effects on protein structure and function in contrast to techniques such as SUnSET (surface sense of translation), which utilize the tRNA analogue puromycin, resulting in truncation of the polypeptide chain (Schmidt *et al*, 2009).

NCAA labelling has been utilized to study the *de novo* proteome in a wide range of animal models, including *Drosophila melanogaster* (Erdmann *et al*, 2015), *Caenorhabditis elegans* (Liang *et al*, 2014), and rodents, and has even been proposed for use in human patients (McBride *et al*, 2014). In mice, it has been used to study how protein synthesis is altered by environmental enrichment (Alvarez-Castelao *et al*, 2019) and impaired in models of frontotemporal dementia (Evans *et al*, 2019). NCAAs like AHA have typically been delivered to the brain via either dietary supplementation (McClatchy *et al*, 2015) or intraperitoneal (i.p.) injection (Evans *et al*, 2019, Evans *et al* 2021b), resulting in long minimum labelling periods of upwards of 16 hours. As such, until now AHA labelling has been unable to be used to robustly study the spatial and temporal dynamics of learning-induced protein synthesis.

Here, we overcome these technical limitations by delivering AHA to the brain via the retro-orbital (R.O.) sinus. We report that retro-orbital injection of AHA into awake mice results in rapid labeling of the *de novo* proteome in time periods as short as 30 minutes. By combining FUNCAT with tissue clearing and light sheet microscopy, we reveal that this new technique rapidly labels the *de novo* proteome throughout the mouse brain. Harnessing this technique, we characterize the spatial and temporal dynamics of protein synthesis induced by training in an auditory threat conditioning paradigm. We found that immediately following training, conditioning-induced protein synthesis occurs predominately in the amygdala, before becoming more pronounced in the hippocampus and somatosensory cortex six hours later. Together, our findings show that NCAA labelling can be used to dissect the spatio-temporal complexities of conditioning-induced protein synthesis in the rodent brain.

## Results

### Rapid labeling of the brain de novo proteome enabled by retro-orbital injection of AHA

We first sought to determine whether retro-orbital delivery improved the temporal resolution of AHA labeling compared to other delivery techniques. To address this issue, we utilized FUNCAT-western blot (FUNCAT-WB) analysis to quantify the amount of *de novo* proteomic labelling in the hippocampal tissue of mice after two hours of labeling. Mice were administered 50 mg/kg of AHA either via intraperitoneal, intravenous (tail-vein), or retro-orbital injection. 50 mg/kg was administered as this was previously shown to be the optimal dosage of AHA when delivered via i.p. injection. A treatment period of two hours was selected as these previous studies that utilized i.p. injection observed very limited AHA labeling at this time.

FUNCAT-WB quantification revealed that with two hours of labeling, retro-orbital injection of AHA resulted in nearly a 4-fold increase in *de novo* proteomic labeling compared to i.p injection, and nearly double the labeling achieved via tail-vein injection (Fig. 1B). We next determined the temporal dynamics of AHA labeling following retro-orbital delivery. Using FUNCAT-WB we were able to detect significant AHA labeling as little as 30 minutes post-injection, with the amount of AHA labeling peaking at eight hours post injection (Fig. 1C). Following this, we determined that 50 mg/kg of AHA is the optimal dosage for retro-orbital injection, with higher doses not increasing the amount of *de novo* proteomic labeling two hours post-injection (Fig. 1D). Finally, we used FUNCAT-IHC to demonstrate that the observed AHA labeling is protein synthesis-dependent as co-injection of the protein synthesis inhibitor cycloheximide (CHX) significantly reduced the FUNCAT signal (Fig. 1E).

### Brain wide visualization of protein synthesis by Fun-DISCO

Immunohistochemistry with FUNCAT labeling in slices is limited by the spatial resolution of individual slices and inherently adds bias to an analysis of region-specific changes in protein synthesis tied to a behavioral task. To spatially resolve region-specific changes in protein synthesis tied to conditioning in unbiased manner we developed Fun-DISCO, which leverages Click-3D (Tamura *et al*, 2024) paired with FUNCAT labeling, brain clearing (Friedmann *et al*, 2020), and light sheet microscopy to visualize the nascent proteome brain-wide (Fig. 2A).

**Figure 2:**
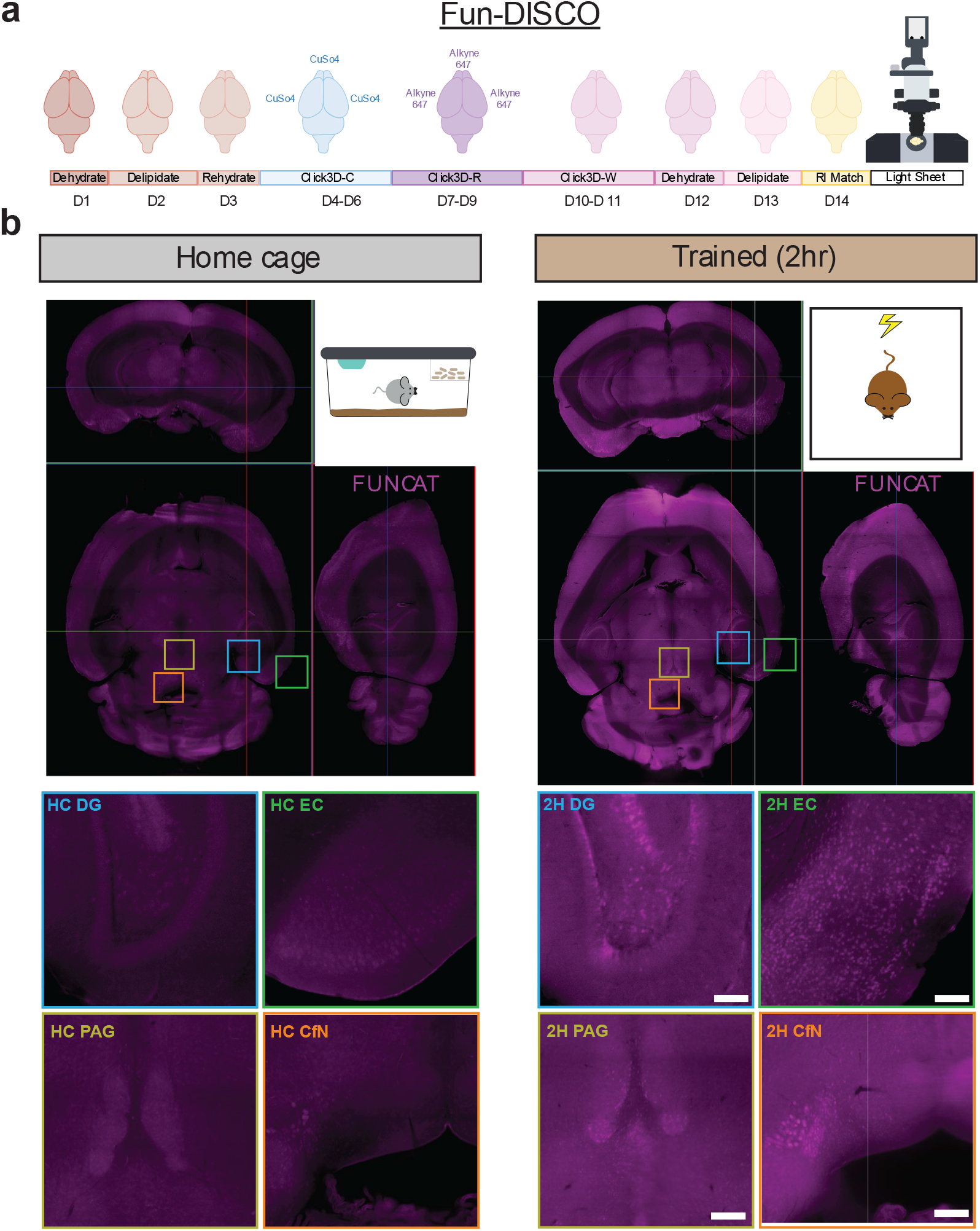
Fun-DISCO enables brain wide visualization of conditioning induced protein synthesis. **(A)** FUNCAT staining was combined with a modified version of iDISCO to visualize protein synthesis in the brains of mice which were administered AHA for two hours via retro-orbital injection, immediately after contextual fear conditioning. **(B)** Compared to home-cage controls, these trained mice appear to exhibit increased FUNCAT staining in brain regions typically associated with contextual memory, such as the dentate gyrus and entorhinal cortex, as well as other brains regions such as the periaqueductal gray (PAG) and cuneiform nucleus (CfN) not typically associated with conditioning induced plasticity. Scale bar = 200μm

As a proof of principle, we conducted FUNCAT-iDISCO (Fun-DISCO) on brains perfused 2 hours after retro-orbital delivery of AHA following a contextual fear conditioning paradigm (Fig. 2). Compared to mice injected with AHA after resting in their home cage, AHA injected conditioned animals showed robust region-specific intensities of protein synthesis including in the dentate gyrus and the entorhinal cortex (Fig. 2B), two regions that have previously been implicated in threat memory formation (Feng *et al*, 2021; Bernier *et al*, 2017). In addition, we noticed increased FUNCAT staining intensity in a number of thalamic and midbrain nuclei in trained mice (Fig. 2B). Thus, Fun-DISCO has the capacity to discover novel regions involved in learning-induced plasticity in an unbiased way that may be missed through conventional sectioning approaches.

### Rapid changes in protein synthesis in the amygdala occur immediately following auditory threat conditioning

Next, we sought to leverage the vast improvement in temporal resolution enabled by the retro-orbital delivery of AHA to explore the spatial and temporal dynamics of learning-induced *de novo* protein synthesis immediately following training in a different behavioral paradigm, namely auditory threat conditioning. Here, mice were trained to associate hearing a tone (conditioned stimulus, CS) with receiving a foot shock (unconditioned stimulus, US), and were compared to mice which did not receive foot shocks, as well as mice where the foot shock and tone were unpaired (Fig. 3). FUNCAT-IHC analysis revealed that neuronal *de novo* protein synthesis was significantly increased in the amygdala in the two hours immediately following conditioning (Fig. 3). This is consistent with previous studies that have shown memory consolidation following this associative conditioning paradigm to be dependent upon neuronal protein synthesis the amygdala (Shrestha *et al*, 2020a). We also observed that hippocampal neuronal protein synthesis was increased in both our paired and unpaired groups compared to non-shock controls, with this increase being significantly higher in the paired mice (Fig. 3). Our analysis also revealed that protein synthesis levels in the somatosensory cortex remained unchanged in the two hours immediately following training (Fig. 3).

**Figure 3:**
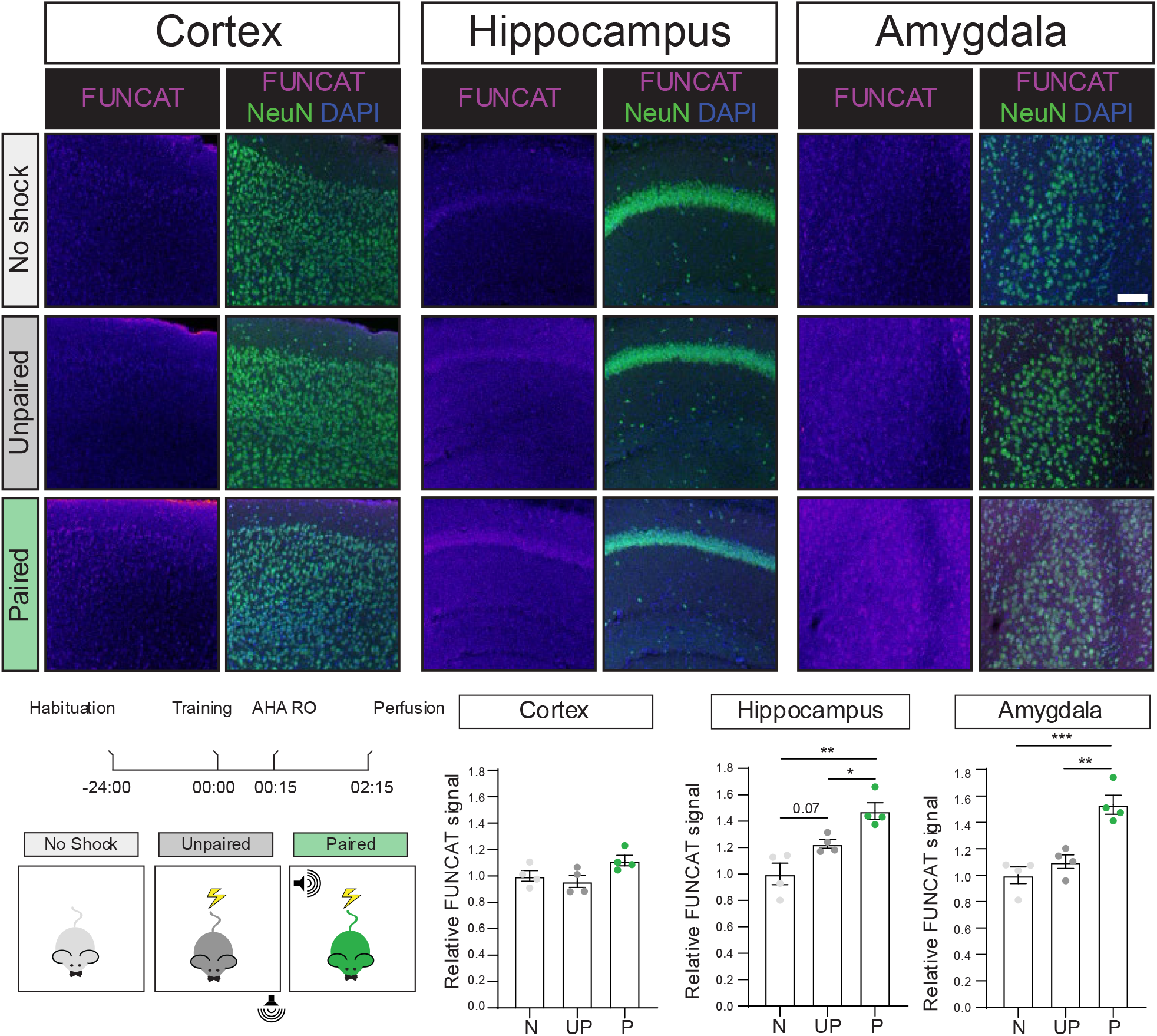
Auditory threat conditioning elevates neuronal protein synthesis in the hippocampus and amygdala immediately following training. Trained mice were conditioned to associate receiving a foot shock (unconditioned stimulus; US) upon hearing an auditory cue (conditioned stimulus; CS) within a specific context. Immediately following training, mice were administered 50 mg/kg AHA via RO injection before being perfused 2 hours later. These mice were compared to an unpaired control group, where mice received the foot shock independently from the CS, as well as non-shock control group, where mice were exposed to the context but not the US or CS. FUNCAT-IHC analysis revealed that neuronal protein synthesis was signifcantly increased in amygdala and the CA1 region of the hippocampus of mice trained via the ACFC compared to non-shock and unpaired controls, with NeuN immunoreactivity being used to identify neurons. FUNCAT signal remained unchanged in the somatosensory cortex (one-way ANOVA, Tukey’s MCT, n=4 animals, ≥3 sections per animal). Scale bar = 100 μm. Error bars = S.E.M. *= p≤ 0.05, **= p≤ 0.01, ***= p≤ 0.001, ****= p≤ 0.001.

### Auditory threat conditioning induces multiple temporally shifted windows of protein synthesis across different brain regions

Given our improvements made to the temporal resolution of *de novo* proteomic labelling *in vivo*, we next determined how learning-induced protein synthesis changes throughout the process of memory consolidation. The consensus in the field is that the neuronal changes associated with memory consolidation occur at different time points, with some changes immediately following training, and others occurring hours or even days later. We therefore chose to examine neuronal protein synthesis between five and seven hours following auditory threat conditioning (Fig. 4). Although FUNCAT-IHC analysis revealed that neuronal protein synthesis in the amygdala and hippocampus was still significantly elevated (Fig. 4), more robust differences were observed in the somatosensory cortex at this later timepoint. Unlike immediately following training, at 5-7 hours post training, both the paired and unpaired mice exhibited a large increase in *de novo* protein synthesis in the layer 2/3 neurons of the somatosensory cortex compared to no shock controls. These findings suggest that *de novo* protein synthesis in this brain region may be involved in later processes of memory consolidation compared to the amygdala and hippocampus.

**Figure 4:**
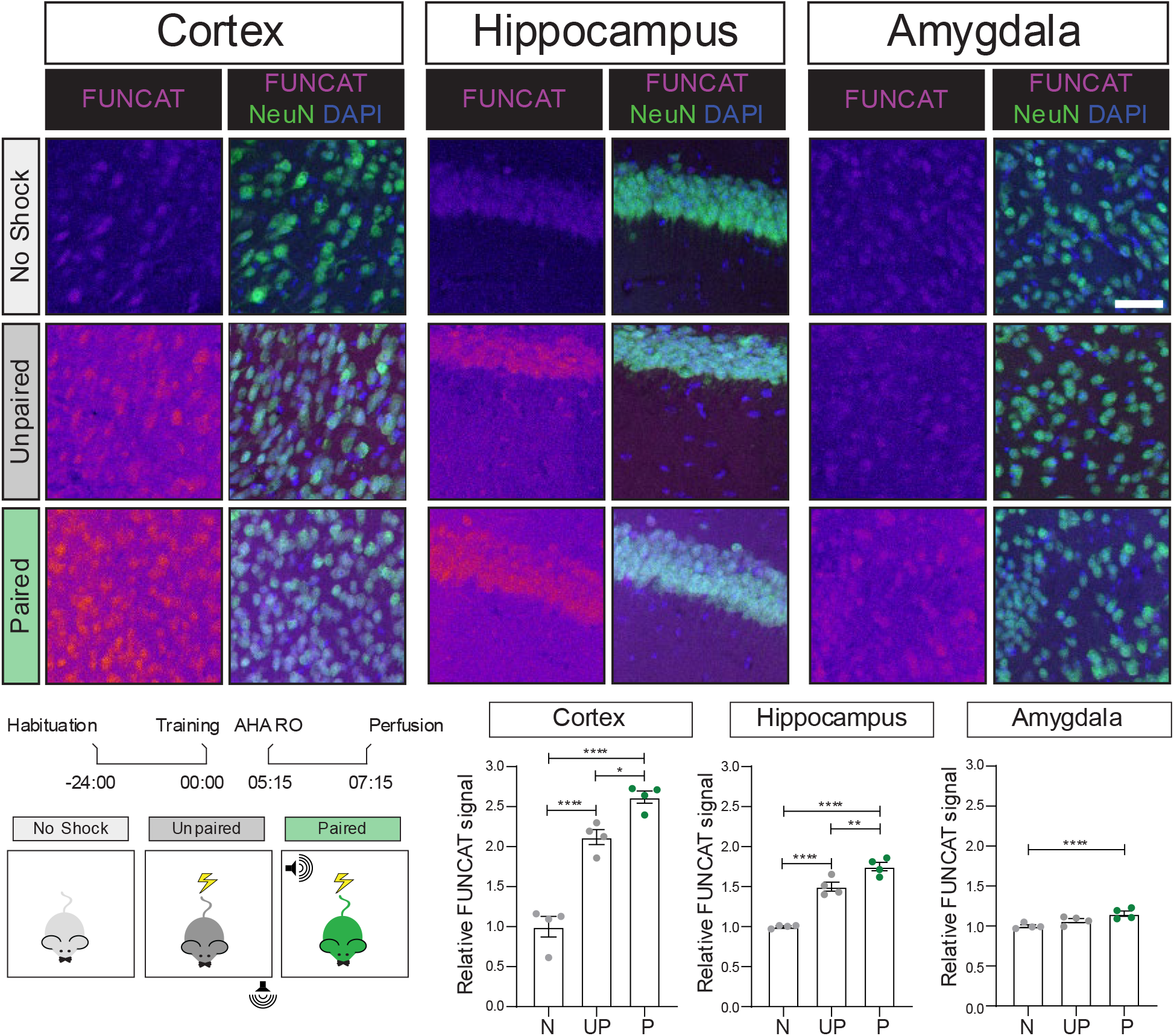
A secondary wave of conditioning-induced protein synthesis occurs in layer 2/3 somatosensory neurons five hours after auditory threat conditioning. Mice in the trained, unpaired, and non-shock groups were administered 50 mg/kg AHA via RO injection 5 hours after their respective behavioral paradigms and perfused 2 hours later. FUNCAT-IHC analysis revealed that neuronal protein synthesis remained significantly elevated in amygdala in trained mice compared to non-shock controls. Interestingly at this later time point, protein synthesis was significantly increased in the CA1 neurons of the hippocampus and layer 2/3 neurons of the somatosensory cortex in both trained and unpaired mice when compared to the non-shock controls, with this effect being more pronounced in trained mice (one-way ANOVA, Tukey’s MCT, n=4 animals, ≥3 sections per animal). Scale bar = 50μm. Error bars = S.E.M. *= p≤ 0.05, **= p≤ 0.01, ***= p≤ 0.001, ****= p≤ 0.001

## Discussion

For over 60 years it has been known that the formation of LTM is dependent upon protein synthesis, but the temporal and spatial landscape of conditioning-induced *de novo* protein synthesis remains relatively underexplored. Herein, through combining the retro-orbital delivery of AHA with Fun-DISCO, we have significantly improved the spatial and temporal resolution of *de novo* proteomic labeling *in vivo*, allowing for the visualization of multiple windows of conditioning-induced protein synthesis throughout the mouse brain for the first time. These advancements enabled us to demonstrate that protein synthesis occurs predominately in the amygdala and hippocampus during the initial stages of auditory threat memory consolidation, before becoming more pronounced in the somatosensory cortex at later time points.

Delivering AHA via retro-orbital injection allowed us to gain greater insights into the spatiotemporal dynamics of conditioning-induced protein synthesis compared to previous studies that have used AHA labeling in the context of memory. In the first of these studies, the authors delivered AHA to mice trained using a modified active place avoidance paradigm, where they observed increased protein synthesis in the hippocampus but not in the somatosensory cortex (Evans *et al*, 2020). However, as this study relied upon i.p. injection, the authors were forced to label the *de novo* proteome over a period of 16 hours. Our findings suggest that this extended labeling period would have prevented the observation of the multiple, distinct windows of conditioning-induced protein synthesis that occur during this timeframe, meaning that FUNCAT signal was only found to be increased in areas of the brain where protein synthesis was constantly upregulated.

A more recent study showed that dietary depletion of methionine can increase AHA labeling efficiency in mice (Sharma *et al*, 2023). The authors in this study used AHA to examine astrocytic protein synthesis in mice following viral-mediated modulation of eIF2α (eukaryotic initiation factor 2α) phosphorylation. By depleting methionine for one week, the authors were able to observe *de novo* protein labelling three hours after i.p. injection of AHA. When designing our approach for examining learning-induced protein synthesis we opted to avoid methionine depletion, reasoning that it may interfere with metabolically sensitive processes involved in memory formation. Methionine is an essential amino acid found in nearly all proteins (Lim *et al*, 2019) and as such any prolonged dietary restriction may lead to hypomethoninemia, which can result in a host of neurological symptoms (Kripps *et al*, 2022). As such, any behavioral task or memory-related readout paired with methionine depletion should be interpreted with caution. This is especially the case in models of neurodegenerative and neurodevelopment disease where metabolism is already impacted (Gruss, 2004; Griffin & Bradshaw, 2017; Cunnane *et al*, 2020).

By improving the temporal specificity of AHA labelling, we were able to reveal that protein synthesis is elevated in the amygdala and hippocampus immediately following auditory threat conditioning and remains elevated during later stages of memory consolidation (Fig. 3&4). These findings add further support to the plethora of evidence demonstrating that protein synthesis in these two brain regions are critically involved in the processing of threat memory, with the amygdala thought to enable long-term association between the administration of the CS (the tone) and the US (the foot shock), whereas the hippocampus is thought to allow long-term association between the foot shock and the spatial context (Phillips & LeDoux, 1992).

Notably, we observed that in the somatosensory cortex, neuronal *de novo* protein synthesis is unchanged immediately following auditory threat conditioning but becomes significantly elevated five hours later (Fig. 3&4). In the auditory threat conditioning paradigm, lesioning of the somatosensory cortex in rats does not prevent the formation of a conditioned response (freezing) to the CS (tone) or the context in which the CS is presented (Phillips & LeDoux, 1992). Instead, during auditory threat conditioning it is thought that these cortical regions are required for the long-term storage of these memories (Bergstrom, 2016). It is possible that the delayed window of neuronal protein synthesis that we observed in the somatosensory cortex may help facilitate this storage.

When seeking to identify the brain regions and cell types in which protein synthesis is required to facilitate memory formation, the field has done so using candidate approaches. In these studies, various pharmacological, genetic, and chemogenetic techniques have been leveraged to block protein synthesis in brain regions and specific cell types already thought to be involved in the particular form of memory being tested (Shrestha *et al*, 2020a, 2020b; Ozawa *et al*, 2017). Here, by combining retro-orbital delivery of AHA with Fun-DISCO, we have enabled the rapid labeling and visualization of the *de novo* proteome throughout the entirety of the mouse brain. This technological advancement will enable researchers to, in a non-biased manner, identify both where and when protein synthesis is altered during the different stages of memory consolidation.

Although the reliance of LTM upon protein synthesis was first described in 1963, researchers are yet to identify which proteins need to be synthesized, in which cells, and at what timepoint in order to facilitate memory consolidation. Here, by significantly advancing the spatial and temporal resolution of AHA labeling *in vivo*, we hope to enable researchers to more accurately map changes in the *de novo* proteomic landscape that underlies long-term changes in complex behaviors.

## Methods

### Animal Care

Wild-type C57/bl6J male and female mice of 3-5 months of age were provided with both food and water *ad libitum* and were maintained on a 12h/12h light/dark cycle at a stable temperature (78°F) and humidity (40–50%). All procedures involving the use of animals were performed in accordance with the guidelines of the National Institutes of Health and were approved by the Institutional Animal Care and Use Committee (IACUC, protocols 221-1143 and 221-1145).

### Contextual and cued-threat conditioning

Rodent behavioral training and analysis was performed during the light cycle of the mice, with an even distribution of male and female mice across groups. For contextual fear conditioning, mice were placed into fear conditioning chamber (Coulbourn instruments) for 270 seconds before receiving a 2 second, 0.5mA foot-shock. This foot-shock (conditioned stimulus-CS) was repeated 100 and 200 seconds later, with the mouse being removed from the chamber 150 seconds after the administration of the final shock, spending a total of 12 minutes in the chamber.

For cued-threat conditioning, all mice were first habituated to the fear conditioning chamber for 15 minutes, one-day prior to training. On day two, trained mice were placed into the chamber for 270 seconds, before being presented with a 5kHz, 85 dB pure tone for 30s, culminating with the administration of a 2 second, 0.5mA foot-shock. This paired-tone presentation (conditioned stimulus-CS) was repeated 100 and 200 seconds later, with the mouse being removed from the chamber 150 seconds after the administration of the final shock, spending a total of 12 minutes in the chamber. For the unpaired controls, mice spent a total of 12 minutes in the chamber. Mice were administered a 2 second, 0.5mA foot-shock 300, 430 and 530 seconds after being placed in the chamber. These mice were presented with a 5kHz, 85 dB pure tone for 30s at 100, 350 and 600 seconds after being placed in the chamber. Non-shock controls were placed in the chamber for a total of 12 minutes and were not administered a foot shock or presented with the audio cue. Freezing behavior was automatically measured by Freeze Frame 4 software (ActiMetrics).

Mice were then administered AHA via RO injection either immediately after being removed from the chamber, or 5 hours later.

### Delivery of azidohomoalanine to awake mice

Azidohomoalanine (Vector Laboratories, CCT-1106) was dissolved in phosphate-buffered saline (PBS) prior to delivery to awake via either intraperitoneal (IP), intravenous (tail-vein), or retro-orbital (RO) injection. A total injection volume of 50uL was used for all three delivery methods. For delivery of AHA via RO injection, a 0.5% proparacaine hydrochloride ophthalmic solution (Covetrus, 2963726) was administered to the eye 2 minutes prior to injection. RO injection in awake mice was then performed by two researchers, with the first restraining the mouse’s head before subsequently drawing back the skin below the eye and second researcher using a 27G needle to inject into the retro-bulbar sinus. For experiments requiring the inhibition of protein synthesis using the protein synthesis inhibitor cycloheximide (CHX, Millipore Sigma, 01810), 10 mg/kg CHX was delivered alongside AHA via RO injection.

Mice were then deeply anesthetized with isoflurane before being intracardially perfused with 20 mL of PBS. For FUCNAT-WB analysis, the hippocampus was then dissected before being snap-frozen. Mice which underwent subsequent FUNCAT/IHC analysis were then intracardially perfused with 20 mL of 4% paraformaldehyde (PFA, ThermoFisher, 50-980-494).

### FUNCAT-Western Blot analysis

Snap-frozen samples were lysed via sonication in radioimmunoprecipitation assay (RIPA) buffer (Cell Signaling, 9806) with Halt protein inhibitor cocktail (ThermoFisher, 78,438) and 1 mM phenylmethylsulfonyl fluoride (ThermoFisher, 36,978), with the EZQ protein quantification assay (Invitrogen, R33200) being used to determine protein concentration.

Newly synthesized proteins were then labelled with IRDye® 800CW Alkyne Infrared Dye (LI-COR, 929-60002) using the Click-&-Go® Protein Reaction Buffer Kit (Vector Laboratories, CCT-1262) as per the manufacturer’s instructions. Briefly, newly synthesized proteins contained within 100 μg of sample were labelled with 20 μM 800CW Alkyne at RT for 30 minutes prior to undergoing chloroform-methanol precipitation as previously described (Evans *et al*, 2024). Samples were then resuspended in 100uL of RIPA buffer with 2.5% SDS, before being denatured by boiling at 95°C for 5 minutes in 1X Laemmli sample buffer (Bio-Rad-1610747) with 5% 2-Mercaptoethanol (Millipore-Sigma, M6250).

Samples were then separated via SDS-PAGE using a 4-20% Tris-Glycine gel (Invitrogen, XP04205BOX) before being transferred to a PVDF membrane (Invitrogen, IB24002) using the iBlot semidry transfer system (Invitrogen, IB2100). The total protein stain REVERT (LI-COR, 926– 10,011) was used for normalization.

### Immunohistochemistry, FUNCAT and Fun-DISCO

For IHC and FUNCAT staining, brains fixed with 4% PFA were submerged in PBS with 30% w/v sucrose for 48 hours prior to being sectioned at 40 μM thickness using a vibratome (Leica, VT1200). Samples were then blocked and permeabilized for 1h at RT under constant agitation in 5% bovine serum albumin, 5% normal goat serum, and 0.5% triton-x in PBS. Newly synthesized proteins were then labelled with 5μM Alexa 647 alkyne (Vector Laboratories, CCT-1301) using the Click-&-Go® Cell Reaction Buffer Kit (Vector Laboratories, CCT-1263) as per the manufacturer’s instructions. Neurons were visualized using a guinea pig IgG anti-NeuN primary antibody (Synaptic Systems, 266 004, 1:1000) and Alexa-488 labeled goat anti-Guinea Pig IgG (H+L) (Invitrogen, A-11073). DAPI was used to stain cell nuclei and samples were mounted in ProLong Gold mounting media (ThermoFisher, P10144).

For Fun-DISCO, brains were transcardially perfused with heparinized (.01%) PBS followed by 4%PFA. Brains were then post-fixed overnight in 4% PFA while rotating on an orbital shaker at 4oC. Following post-fixation, brains were washed 3x (for 2hrs, 4hrs, and overnight) in PBS and transferred to 5mL Eppendorf tubes. Brains were dehydrated in increasing concentrations MeOH (20%, 40%, 60%, and 80% 1hr each) diluted in B1N solution without sodium azide (2% glycine, .1% tritonX-100, and .01% 10N NaOH) at room temperature (RT) on an orbital shaker. Brains were then further dehydrated 4x in 100% MeOH for 1hr at RT. To delipidate the samples, a 2:1 solution of dichloromethane (DCM, Sigma 270997) and MeOH overnight (ON). The next day brains were further depilated in 100% DCM 3x for 1hr at RT with shaking. Samples were then washed 2x 1hr in 100% MeOH followed by rehydration in a MeOH/B1n series (60%, 40%, 20%) for 1 hr each at RT with shaking, followed by 100% B1N ON. The next day samples were begun on an adapted Click 3D protocol (Tamura et al., 2024): Brains were incubated in 4mL of Click3D-C solution (10mM Hepes (pH 7.3), 900mM NaCl, 10% w/v DMSO, 4mM THPTA (Vector Labs CCT-1010, 2mM CuSO4 added to sterile H2O with vertexing after each addition) for 2 days at 370 on a 3600 vertical rotator. The Click3D-C solution was exchanged with new Click3D-C and rotated ON at 370 on a 3600 vertical rotator. The next day Click3D-C solution was exchanged for Click3D-R solution (10mM Hepes, 900mM NaCl, 10% w/v DMSO, 2mM CuSO4, 15 μM Alexa 647 alkyne (Vector Laboratories, CCT-1301), and freshly prepared 100mM NaAsc (Sigma, PHR1279), added to sterile H2O and vortexed after each addition) and rotated 2x ON at 370 on a 3600 vertical rotator with a fresh exchange after the first night. The Click3D-R solution was exchanged 1 more time and incubated at 370 on a 3600 vertical rotator for 2 hrs. Then the solution was exchanged for Click3D-W (0.1M EDTA (pH 8), .2% Triton x-100 in sterile H2O) and incubated for 2 hrs 4x at 370 on a 3600 vertical rotation with an exchange of solution after each period followed by 1x incubation ON. The following day the brains were washed 3x 1hr in PBS followed by dehydration in a MeOH series (20%, 40%, 60%, 80%, 100%, 100%, 100%) 1hr each diluted in H2O on an orbital shaker at RT. The brains were further delipidated ON in a 2:1 DCM/MeOH ON at RT on an orbital shaker. The next day the brains were incubated 2x 1hr in 100% DCM. Finally the brains were transferred to a 5mL amber borosilicate vial filled with benzyl ether (Sigma, 108014) for refractive index matching and stored until imaging.

### Imaging and image analysis

For FUNCAT-Western blot analysis, membranes were imaged using a LI-COR Odessey M Scanner with the LI-COR Emperia Studio software being used for quantification.

For FUNCAT-IHC analysis, 15 μm thick Z-stack images were taken using a Leica SP8 Confocal microscope with maximum intensity projections being created in ImageJ. Image analysis was performed blinded in ImageJ with NeuN immunoreactivity being used to generate a mask. Mean gray value was then measured within this neuronal mask for each image, with no significant difference being detected between the area of these masks across groups. A minimum of 3 sections per animal were analyzed, with each data point representing an average of these sections. For Fun-DISCO, cleared whole brains were imaged in benzyl ether using a Zeiss Z1 light sheet microscope at 5× magnification with the Zeiss Zen Black software. Image stitching and generation of orthogonal views were performed using Zeiss Zen Blue.

### Statistical analysis

GraphPad Prism 10.1.2 was used for statistical analysis, with a one-way ANOVA with Tukey’s multiple comparison test (MCT), or Student’s T.Test being used as appropriate. All values are given as mean ± standard error of the mean. Significance was defined as *p < 0.05, **p < 0.01, ***p < 0.001, ****p < 0.0001.

## Acknowledgement

The authors would like to acknowledge the contributions Maggie Donohue and Jenesha Rawlani to this work.

## References

Alvarez-Castelao B, Schanzenbächer CT, Langer JD & Schuman EM (2019) Cell-type-specific metabolic labeling, detection and identification of nascent proteomes in vivo. Nat Protoc 14: 556–575

Bergstrom HC (2016) The neurocircuitry of remote cued fear memory. Neurosci Biobehav Rev 71: 409–417 doi:10.1016/j.neubiorev.2016.09.028 [PREPRINT]

Bernier BE, Lacagnina AF, Ayoub A, Shue F, Zemelman B V., Krasne FB & Drew MR (2017) Dentate Gyrus Contributes to Retrieval as well as Encoding: Evidence from Context Fear Conditioning, Recall, and Extinction. The Journal of Neuroscience 37: 6359–6371

Carlisle AK, Götz J & Bodea LG (2023) Three methods for examining the de novo proteome of microglia using BONCAT bioorthogonal labeling and FUNCAT click chemistry. STAR Protoc 4

Cunnane SC, Trushina E, Morland C, Prigione A, Casadesus G, Andrews ZB, Beal MF, Bergersen LH, Brinton RD, de la Monte S, et al (2020) Brain energy rescue: an emerging therapeutic concept for neurodegenerative disorders of ageing. Nat Rev Drug Discov 19: 609–633

Dieterich DC, Link AJ, Graumann J, Tirrell DA & Schuman EM (2006) Selective identification of newly synthesized proteins in mammalian cells using bioorthogonal noncanonical amino acid tagging (BONCAT). Proceedings of the National Academy of Sciences 103: 9482– 9487

Elder MK, Erdjument-Bromage H, Oliveira MM, Mamcarz M, Neubert TA & Klann E (2021) Age-dependent shift in the de novo proteome accompanies pathogenesis in an Alzheimer’s disease mouse model. Commun Biol 4: 823

Erdmann I, Marter K, Kobler O, Niehues S, Abele J, Müller A, Bussmann J, Storkebaum E, Ziv T, Thomas U, et al (2015) Cell-selective labelling of proteomes in Drosophila melanogaster. Nat Commun 6: 7521

Evans HT, Benetatos J, van Roijen M, Bodea L-G & Götz J (2019) Decreased synthesis of ribosomal proteins in tauopathy revealed by non-canonical amino acid labelling. EMBO J 38: e101174

Evans HT, Blackmore D, Götz J & Bodea LG (2021a) De novo proteomic methods for examining the molecular mechanisms underpinning long-term memory. Brain Res Bull 169: 94–103

Evans HT, Bodea L & Götz J (2020) Cell-specific non-canonical amino acid labelling identifies changes in the de novo proteome during memory formation. Elife 9: 1–19

Evans HT, Ko T, Oliveira MM, Yu A, Kalavai S V., Golhan EN, Polavarapu A, Balamoti E, Wu V, Klann E, et al (2024) Light-Activatable, Cell-Type Specific Labeling of the Nascent Proteome. ACS Chem Neurosci 15: 3473–3481

Evans HT, Taylor D, Kneynsberg A, Bodea L-G & Götz J (2021b) Altered ribosomal function and protein synthesis caused by tau. Acta Neuropathol Commun 9: 110

Feng H, Su J, Fang W, Chen X & He J (2021) The entorhinal cortex modulates trace fear memory formation and neuroplasticity in the mouse lateral amygdala via cholecystokinin. Elife 10

Flexner JB, Flexner LB & Stellar E (1963) Memory in Mice as Affected by Intracerebral Puromycin. Science (1979) 141: 57–59

Fonseca R, Nägerl UV & Bonhoeffer T (2006) Neuronal activity determines the protein synthesis dependence of long-term potentiation. Nat Neurosci 9: 478–480

Friedmann D, Pun A, Adams EL, Lui JH, Kebschull JM, Grutzner SM, Castagnola C, Tessier-Lavigne M & Luo L (2020) Mapping mesoscale axonal projections in the mouse brain using a 3D convolutional network. Proceedings of the National Academy of Sciences 117: 11068–11075

Griffin JWD & Bradshaw PC (2017) Amino Acid Catabolism in Alzheimer’s Disease Brain: Friend or Foe? Oxid Med Cell Longev 2017

Gruss M (2004) Age- and region-specific imbalances of basal amino acids and monoamine metabolism in limbic regions of female Fmr1 knock-out mice. Neurochem Int 45: 81–88

Hinz F, Dieterich D & Schuman E (2013) Teaching old NCATs new tricks: using non-canonical amino acid tagging to study neuronal plasticity. Curr Opin Chem Biol 17: 738–746

Koren SA, Hamm MJ, Meier SE, Weiss BE, Nation GK, Chishti EA, Arango JP, Chen J, Zhu H, Blalock EM, et al (2019) Tau drives translational selectivity by interacting with ribosomal proteins. Acta Neuropathol 137: 571–583

Kripps KA, Sremba L, Larson AA, Van Hove JLK, Nguyen H, Wright EL, Mirsky DM, Watkins D, Rosenblatt DS, Ketteridge D, et al (2022) Methionine synthase deficiency: Variable clinical presentation and benefit of early diagnosis and treatment. J Inherit Metab Dis 45: 157–168

Liang V, Ullrich M, Lam H, Chew YL, Banister S, Song X, Zaw T, Kassiou M, Götz J & Nicholas HR (2014) Altered proteostasis in aging and heat shock response in C. elegans revealed by analysis of the global and de novo synthesized proteome. Cellular and Molecular Life Sciences 71: 3339–3361

Lim JM, Kim G & Levine RL (2019) Methionine in Proteins: It’s Not Just for Protein Initiation Anymore. Neurochem Res 44: 247–257

Lima RH, Rossato JI, Furini CR, Bevilaqua LR, Izquierdo I & Cammarota M (2009) Infusion of protein synthesis inhibitors in the entorhinal cortex blocks consolidation but not reconsolidation of object recognition memory. Neurobiol Learn Mem 91: 466–472

McBride WJ, Souza CDA & Goldenberg DM (2014) In vivo copper-free click chemistry for delivery of therapeutic and/or diagnostic agents.

McClatchy DB, Ma Y, Liu C, Stein BD, Martínez-Bartolomé S, Vasquez D, Hellberg K, Shaw RJ & Yates JR (2015) Pulsed Azidohomoalanine Labeling in Mammals (PALM) Detects Changes in Liver-Specific LKB1 Knockout Mice. J Proteome Res 14: 4815–4822

McClatchy DB, Martínez-Bartolomé S, Gao Y, Lavallée-Adam M & Yates JR (2020) Quantitative analysis of global protein stability rates in tissues. Sci Rep 10: 15983

Mohamed MS & Klann E (2023) Autism- and epilepsy-associated EEF1A2 mutations lead to translational dysfunction and altered actin bundling. Proceedings of the National Academy of Sciences 120

Moncada D & Viola H (2007) Induction of Long-Term Memory by Exposure to Novelty Requires Protein Synthesis: Evidence for a Behavioral Tagging. Journal of Neuroscience 27: 7476– 7481

Oliveira MM, Lourenco M V., Longo F, Kasica NP, Yang W, Ureta G, Ferreira DDP, Mendonça PHJ, Bernales S, Ma T, et al (2021) Correction of eIF2-dependent defects in brain protein synthesis, synaptic plasticity, and memory in mouse models of Alzheimer’s disease. Sci Signal 14: eabc5429

Ozawa T, Yamada K & Ichitani Y (2017) Differential requirements of hippocampal de novo protein and mRNA synthesis in two long-term spatial memory tests: Spontaneous place recognition and delay-interposed radial maze performance in rats. PLoS One 12: e0171629

Pedroza-Llinás R, Ramírez-Lugo L, Guzmán-Ramos K, Zavala-Vega S & Bermúdez-Rattoni F (2009) Safe taste memory consolidation is disrupted by a protein synthesis inhibitor in the nucleus accumbens shell. Neurobiol Learn Mem 92: 45–52

Phillips RG & LeDoux JE (1992) Differential contribution of amygdala and hippocampus to cued and contextual fear conditioning. Behavioral Neuroscience 106: 274–285

Schmidt EK, Clavarino G, Ceppi M & Pierre P (2009) SUnSET, a nonradioactive method to monitor protein synthesis. Nat Methods 6: 275–277

Sharma V, Oliveira MM, Sood R, Khlaifia A, Lou D, Hooshmandi M, Hung T-Y, Mahmood N, Reeves M, Ho-Tieng D, et al (2023) mRNA translation in astrocytes controls hippocampal long-term synaptic plasticity and memory. Proceedings of the National Academy of Sciences 120

Shrestha P, Ayata P, Herrero-Vidal P, Longo F, Gastone A, LeDoux JE, Heintz N & Klann E (2020a) Cell-type-specific drug-inducible protein synthesis inhibition demonstrates that memory consolidation requires rapid neuronal translation. Nat Neurosci 23: 281–292

Shrestha P & Klann E (2022) Spatiotemporally resolved protein synthesis as a molecular framework for memory consolidation. Trends Neurosci 45: 297–311

Shrestha P, Shan Z, Mamcarz M, Ruiz KSA, Zerihoun AT, Juan C-Y, Herrero-Vidal PM, Pelletier J, Heintz N & Klann E (2020b) Amygdala inhibitory neurons as loci for translation in emotional memories. Nature 586: 407–411

Tamura I, Sakamoto DM, Yi B, Saito Y, Yamada N, Morimoto J, Takakusagi Y, Kuroda M, Kubota SI, Yatabe H, et al (2024) Click3D: Click reaction across deep tissues for whole-organ 3D fluorescence imaging. Sci Adv 10: 8471

Younts TJ, Monday HR, Dudok B, Klein ME, Jordan BA, Katona I & Castillo PE (2016) Presynaptic Protein Synthesis Is Required for Long-Term Plasticity of GABA Release. Neuron 92: 479–492

